# Screening of Botanical Drugs against SARS-CoV-2 Entry

**DOI:** 10.1101/2021.06.03.447021

**Authors:** Junyuan Cao, Yang Liu, Minmin Zhou, Siqi Dong, Xiaoying Jia, Xiaohao Lan, Yueli Zhang, Jiao Guo, Gengfu Xiao, Wei Wang

**Author notes:** Address correspondence to Wei Wang.

## Abstract

An escalating pandemic caused by the novel severe acute respiratory syndrome coronavirus 2 (SARS-CoV-2) is impacting global health. Specific treatment options for diseases caused by SARS-CoV-2 are largely lacking. Herein, we used a pseudotype virus (pv) bearing the SARS-CoV-2 S glycoprotein to screen a botanical drug library to identify an agent against SARS-CoV-2 entry. All the four hits, including angeloylgomisin O, schisandrin B, procyanidin, and oleanonic acid, were identified for effective inhibition of SARS-CoV-2 S pv entry in the micromolar range. A mechanistic study revealed that these four agents inhibit SARS-CoV-2 S pv entry by blocking S-mediated membrane fusion. Furthermore, angeloylgomisin O, schisandrin B, and oleanonic acid inhibited authentic SARS-CoV-2 with a high selective index (SI). We also showed that all the four hits could also inhibit the entry of pv of Middle East respiratory syndrome coronavirus (MERS-CoV) and newly emerged SARS-CoV-2 variants (D614G, K417N/E484K/N501Y/D614G). In drug combination studies performed in cellular antiviral assays, angeloylgomisin O and schisandrin B displayed synergistic effects in combination with remdesivir. These results indicated that angeloylgomisin O, schisandrin B, procyanidin, and oleanonic acid can inhibit SARS-CoV-2 and that they are potential therapeutic agents for COVID-19.

## INTRODUCTION

COVID-19, caused by severe acute respiratory syndrome coronavirus-2 (SARS-CoV-2), poses a severe threat to global public health as well as economic and social stability, requiring rapid development of treatment and preventive measures (1).

Coronavirus is an enveloped virus with four structural proteins: spike (S), membrane (M), envelope (E), and nucleocapsid (N) proteins. SARS-CoV-2 shares a high degree of sequence identity with SARS-CoV that emerged previously and exploits the same human cell receptor, angiotensin-converting enzyme 2 (ACE2), for infection (2, 3). After the receptor binding domain (RBD) in the S1 subunit of the S protein on the virion binds to the ACE2 receptor on the target cell, the heptapeptide repeat 1 (HR1) and 2 (HR2) domains in the S2 subunit of the S protein interact with each other to form a six-helix bundle (6-HB) fusion nucleus. This interaction brings the virus membrane and cell membrane close together for fusion and infection (4, 5).

SARS-CoV-2 culture and assays need to be carried out in a biosafety level (BSL)-3 laboratory. The vesicular stomatitis virus (VSV) reverse genetics system can provide a safe, reliable, and stable platform to study SARS-CoV-2 S-glycoprotein inhibition by antibodies or small molecules with high assay throughput. Researchers have employed related viral systems that afford virus entry studies at BSL-2 and facilitate rapid inhibitor screening using fluorescence or luminescence-based reporters (6).

More than a year after its emergence, SARS-CoV-2 continues to cause a fervor with identification of new variants hailing from the United Kingdom, South Africa, Brazil, the United States, and India (7–9). The G614 variant was first detected in early 2020 and contains a D614G substitution in its S protein’s RBD, which appears to increase viral transmissibility (10–12). As other new variants emerged over the past few months, the major focus of research has been on identifying further RBD substitutions and their impact on infection. Three other mutations in RBD, K417T/N, E484K, and N501Y, have been identified that affect host cell interactions and susceptibility to antibody-mediated neutralization (13–15).

Natural compounds have been universally used to prevent and cure various illnesses in many countries since ancient times. With the isolation and characterization of the structures of select natural products, these products have become a critical contributor to pharmaceuticals, and several critical pharmaceuticals have been developed that are modeled after natural products (16, 17).

This study focuses on screening drugs targeting the entry step of SARS-CoV-2 infection to block the early stages of viral infection and spread. Using SARS-CoV-2 S VSV-based-pseudovirus (SARS-CoV-2 S pv), we identified that angeloylgomisin O, schisandrin B, procyanidin, and oleanonic acid could inhibit SARS-CoV-2 entry by inhibiting membrane fusion. Our findings may offer new therapeutic strategies for the treatment of COVID-19.

## RESULTS

### Construction of SARS-CoV-2 S pv (ct19)

To perform high-throughput screening under BSL-2 conditions, we constructed a SARS-CoV-2 S pv based on a VSV backbone. The cytoplasmic tail of the S glycoproteins SARS-CoV-2 is highly similar to that of SARS-CoV and carries signals for their retention in the endoplasmic reticulum (ER). Previous studies found that a 19 amino acid deletion in the cytoplasmic tail of SARS-CoV or SARS-CoV-2 S glycoprotein increased the infectivity of the single-cycle pseudotype virus (18–21).

We generated two recombinant plasmids expressing either the wild type (wt) S glycoprotein, pcDNA3.1-SARS-CoV-2 S, or the truncated S glycoprotein missing 19 amino acids from its C-terminus, pcDNA3.1-SARS-CoV-2 S (ct19), which were used to generate SARS-CoV-2 S pv (wt) or the shortened S glycoprotein, SARS-CoV-2 S pv (ct19), respectively.

As shown in Fig. 1A, the *Renilla* luciferase (Rluc) activity of the cells infected with SARS-CoV-2 S pv (ct19) was higher than that of cells infected with SARS-CoV-2 S pv (wt). Next, the VSV copy number was assayed in the different pv harvests, and a higher packaging efficiency was detected in the SARS-CoV-2 S pv (ct19) system than in SARS-CoV-2 S pv (wt) (Fig. 1A), reaching 1.15×10^9^ copies/ml and 2.3×10^8^ copies/ml, respectively. Immunoblots confirmed the expression and accumulation of SARS-CoV-2 S protein in SARS-CoV-2 S pv (ct19) (Fig. 1B), enabling us to use SARS-CoV-2 S pv (ct19) for high-throughput screening (HTS) of potential botanical drug candidates.

**Fig. 1.**
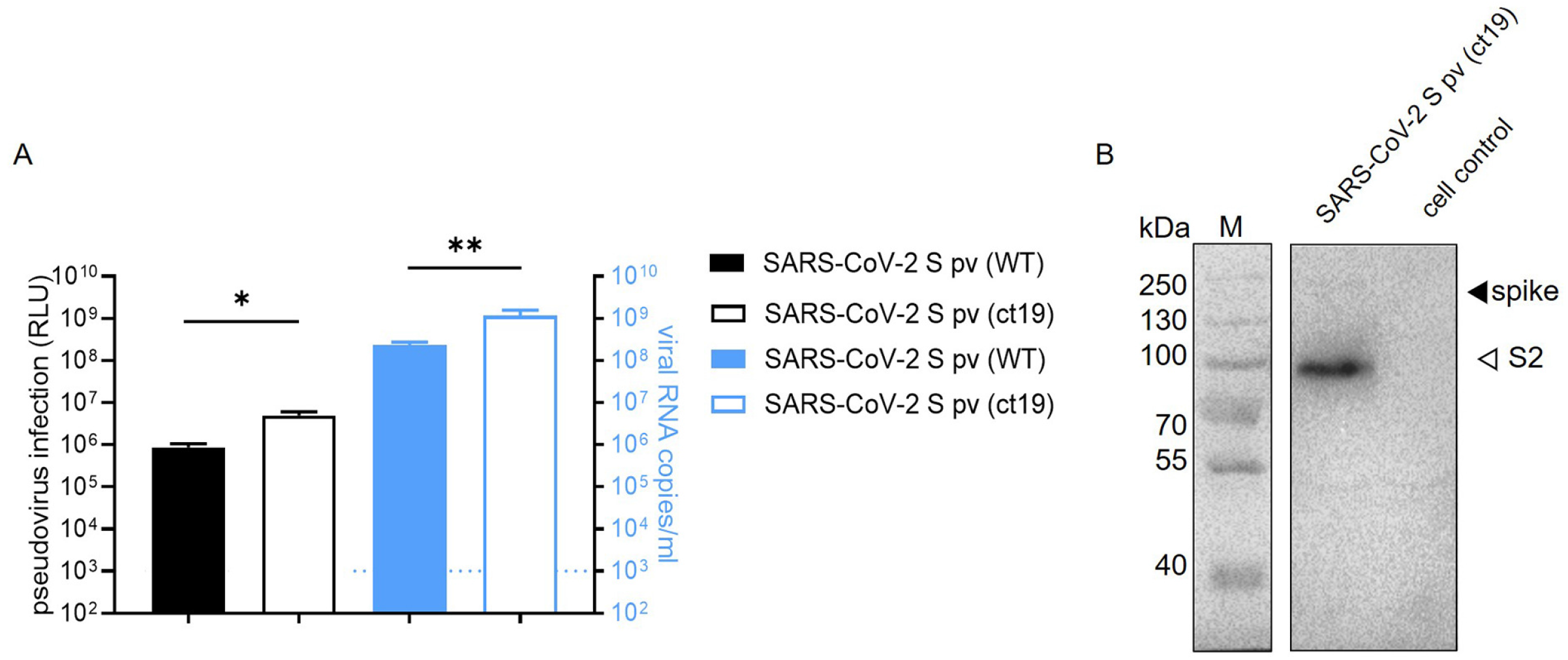
Construction of SARS-CoV-2 S pv (ct19). (A) The left-axis shows the RLU (relative light unit) value detected 24 h after pseudovirus SARS-CoV-2 S pv (ct19) and SARS-CoV-2 S pv (wt) infection. The right y-axis shows virus copy number assays of both viral particles. (B) Immunoblots verifying the incorporation of SARS-CoV-2 spike protein in the pseudovirus. The cell control was the culture supernatant of 293T cell transfected with pcDNA3.1 48-h and infected with VSV pv.

### SARS-CoV-2 S pv (ct19) entry inhibitor screening

The HTS assay conditions were optimized to a seeding Caco-2 cell density of 2.4×10^4^ cells and SARS-CoV-2 S pv (ct19) infective dose of MOI=5 per well in 96-well plates. Under these optimized conditions, the signal-to-basal (S/B) ratio, coefficient of variation, and Z’ factor were 1286.09, 7.35%, and 0.705, respectively, making this assay promising for large-scale inhibitor screening. As shown in Fig. 2A, the HTS assay was performed by screening a library containing 1,037 botanical drugs. Compounds with inhibition of >80% and no apparent cytotoxicity at a concentration of 50 μM were defined as prime candidates. Fifty-eight (5.59%) prime candidates (Fig. 2A and 2B) were selected. These prime candidates were then counter-screened to rule out the inhibition of VSV genome replication and Rluc activity. Four candidates (0.38%), angeloylgomisin O, schisandrin B, procyanidin, and oleanonic acid passed this secondary screen with mild inhibition against VSV (16.85%, 33.65%, 21.02%, and 32.41%, respectively) at 50 μM (Fig. 2C).

**Fig. 2.**
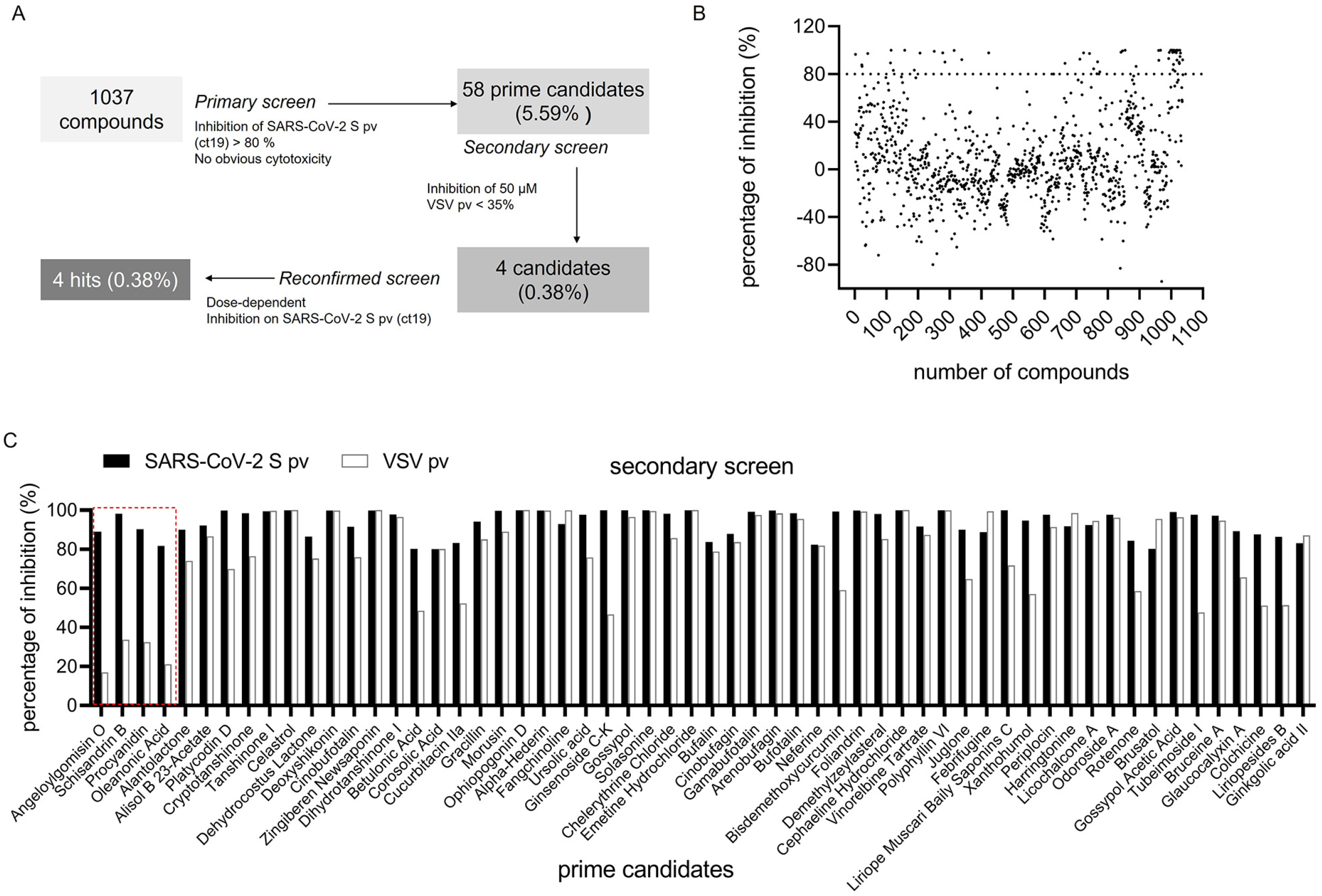
HTS for inhibitors of SARS-CoV-2 entry from a botanical drug library. (A) HTS assay flowchart. (B) HTS of a library of 1,037 natural extracts for primary candidates inhibiting SARS-CoV-2 S pv (ct19) infection. Each dot represents the percent inhibition achieved by each compound at a concentration of 50 μM. (C) Inhibition of 58 candidates against SARS-CoV-2 S pv entry >80%. The second screen of the 58 selected compounds. VSV pv was used as a control to exclude compounds targeting the backbone. The hits depicted in red box showed mild inhibition of VSV pv infection and were designated hits.

### Four hits inhibited SARS-CoV-2 S pv (ct19)

The dose-dependent inhibition of the four hits was further investigated using Caco-2 cells. As shown in Fig. 3, all the four hits suppressed SARS-CoV-2 S pv (ct19) infection in a dose-dependent manner. Angeloylgomisin O inhibited SARS-CoV-2 S pv (ct19) by 90% at 100 μM and had little effect on VSV pv, demonstrating its specific inhibition of SARS-CoV-2 entry (Fig. 3A). Meanwhile, the cell viability was >80% when used at an extremely high concentration (400 μM). Schisandrin B inhibited SARS-CoV-2 S pv (ct19) by 90% at 100 μM, whereas the cell viability was approximately 80% at 400 μM (Fig. 3B). Procyanidin at 50 μM and 100 μM inhibited SARS-CoV-2 S pv (ct19) by approximately 80% and 95%, respectively. The cell viability was approximately 80% at 400 μM (Fig. 3C). Oleanolic acid (50 μM) inhibited SARS-CoV-2 S pv (ct19) by approximately 70% (Fig. 3D). While it is relatively toxic to Caco-2 cells with a 50% cytotoxic concentration (CC_50_) of 53.88 μM. To validate the antiviral effects, four hits were purchased from other commercial sources and tested. The antiviral effects were similar to those observed in our primary screening.

**Fig. 3.**
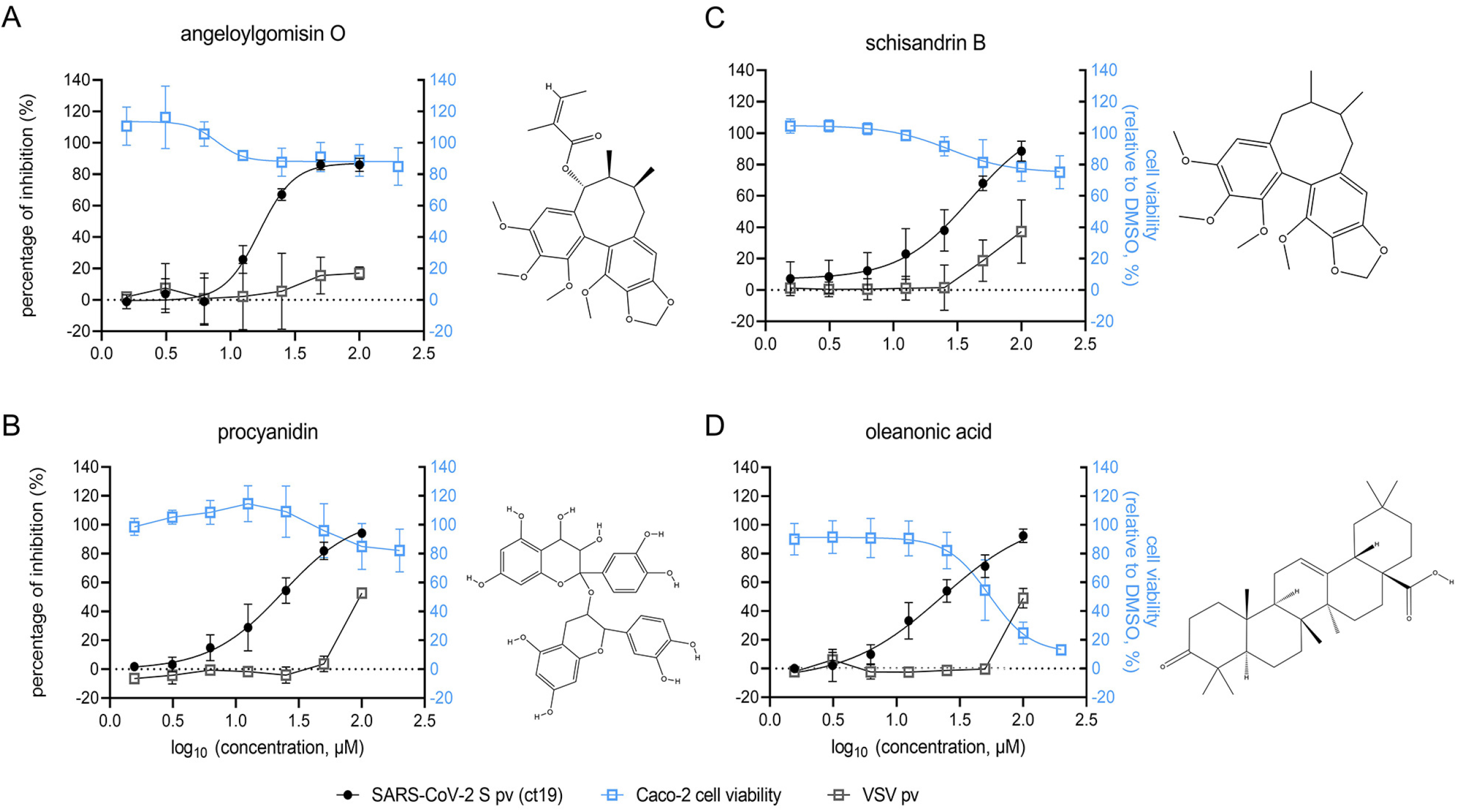
Dose-response curves and the structures of the four hits: angeloylgomisin O (A), schisandrin B (B), procyanidin (C), oleanonic acid (D) for inhibiting SARS-CoV-2 S pv (ct19) infection. (Right) The structure of each hit. Caco-2 cells were seeded at a density of 2.4 × 10^4^ cells per well in 96-well plates. After overnight incubation, cell monolayers were treated in duplicate with the hits at a final concentration; 1 h later, cells were infected with different pv, and the supernatant was removed 1 h post infection. Cell viability was evaluated using MTT assay. Hits at the indicated concentrations was added to pre-seeded Caco-2 cells in 96-well plates. Twenty-four hours later, cell viability was measured. Data are represented as the mean ± standard deviation (SD) from three to four experiments.

### Four hits inhibited SARS-CoV-2 S (ct19)-mediated membrane fusion

To explore the antiviral mechanical properties of the four hits, the effects of the four hits on SARS-CoV-2 S-mediated membrane fusion were examined. Recent research has identified that SARS-CoV-2 induces syncytia formation in the lungs of patients with COVID-19 (22). It was reported that cells infected with SARS-CoV-2 exhibited a typical syncytium phenomenon (5). In this assay, fusion activity was conducted by co-transfecting SARS-CoV-2 S (ct19) and green fluorescent protein (GFP) into Vero E6 cells. After 24 h, the S protein of SARS-CoV-2 induced cell-cell fusion, resulting in the formation of syncytia (Fig. 4A). 25-hydrocholesterol (25HC) was used as the positive control (23, 24). As shown in Fig. 4A, 25 μM of 25HC blocked syncytia formation. Likewise, all the four hits induced a dose-dependent reduction in syncytium size (Fig. 4B). Angeloylgomisin O blocked syncytia formation at a concentration of 6.25 μM, while procyanidin, oleanonic acid, and schisandrin B required higher concentrations (12.5 μM to 25 μM) to achieve the inhibition (Fig. 4B). These results suggested that angeloylgomisin O, schisandrin B, procyanidin, and oleanonic acid could inhibit SARS-CoV-2 entry by inhibiting membrane fusion.

**Fig. 4.**
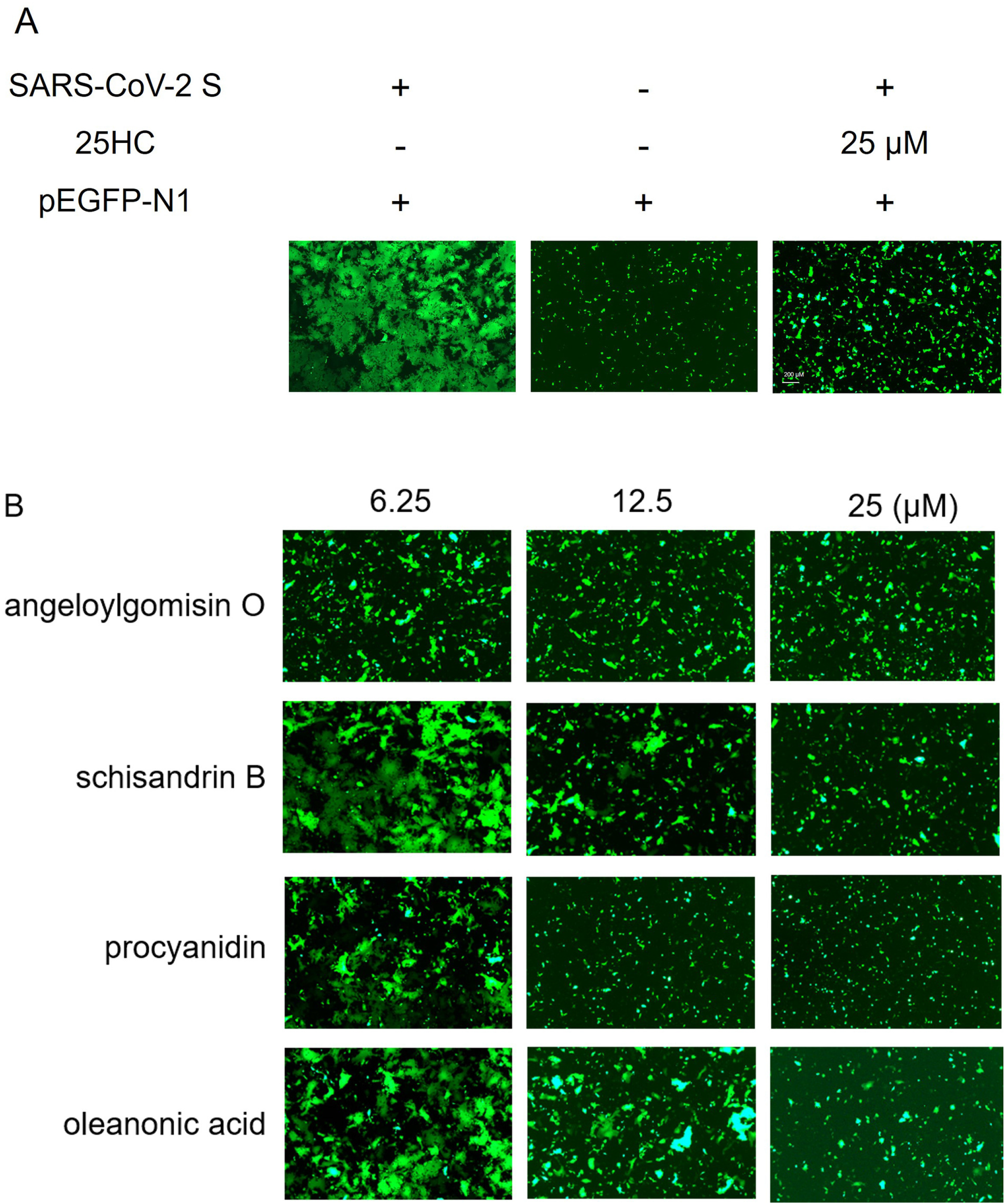
Four hits inhibited SARS-CoV-2 S (ct19)-mediated membrane fusion. (A) SARS-CoV-2 S (ct19)-mediated Cell-cell fusion on Vero E6 cells. (Left) Vero E6 cells that were co-transfected with SARS-CoV-2 S (ct19) and GFP plasmid. (Middle) Cells were transfected with GFP. (Right) Cells were co-transfected with SARS-CoV-2 S (ct19), GFP and treated with 25 μM 25HC. (B) Four hits inhibited SARS-CoV-2 S (ct19)-mediated membrane fusion in dose-depended manner. Vero E6 cells were co-transfected with SARS-CoV-2 S (ct19) and GFP later treated with the four hits in different concentrations. Syncytium formation was visualized 24–36 h later using fluorescent microcopy. Images are representative fields from three independent experiments.

### Four botanical hits inhibited authentic SARS-CoV-2 infection

To evaluate the inhibition of four hits on authentic SARS-CoV-2, an immunofluorescence assay (IFA) was carried out using SARS-CoV-2 NP antibody. As shown in Fig. 5, all the four hits inhibited authentic SARS-CoV-2 in a dose-dependent manner. The IC_50_ values of angeloylgomisin O (3.727 μM), schisandrin B (7.337 μM), and oleanonic acid (1.414 μM) were less than 10 μM, While the IC_50_ value of procyanidin was 32.59 μM. It seemed that angeloylgomisin O, schisandrin B, and oleanonic acid exerted stronger inhibition on the authentic SARS-CoV-2 than on the SARS-CoV-2 S pv (ct19).

**Fig. 5.**
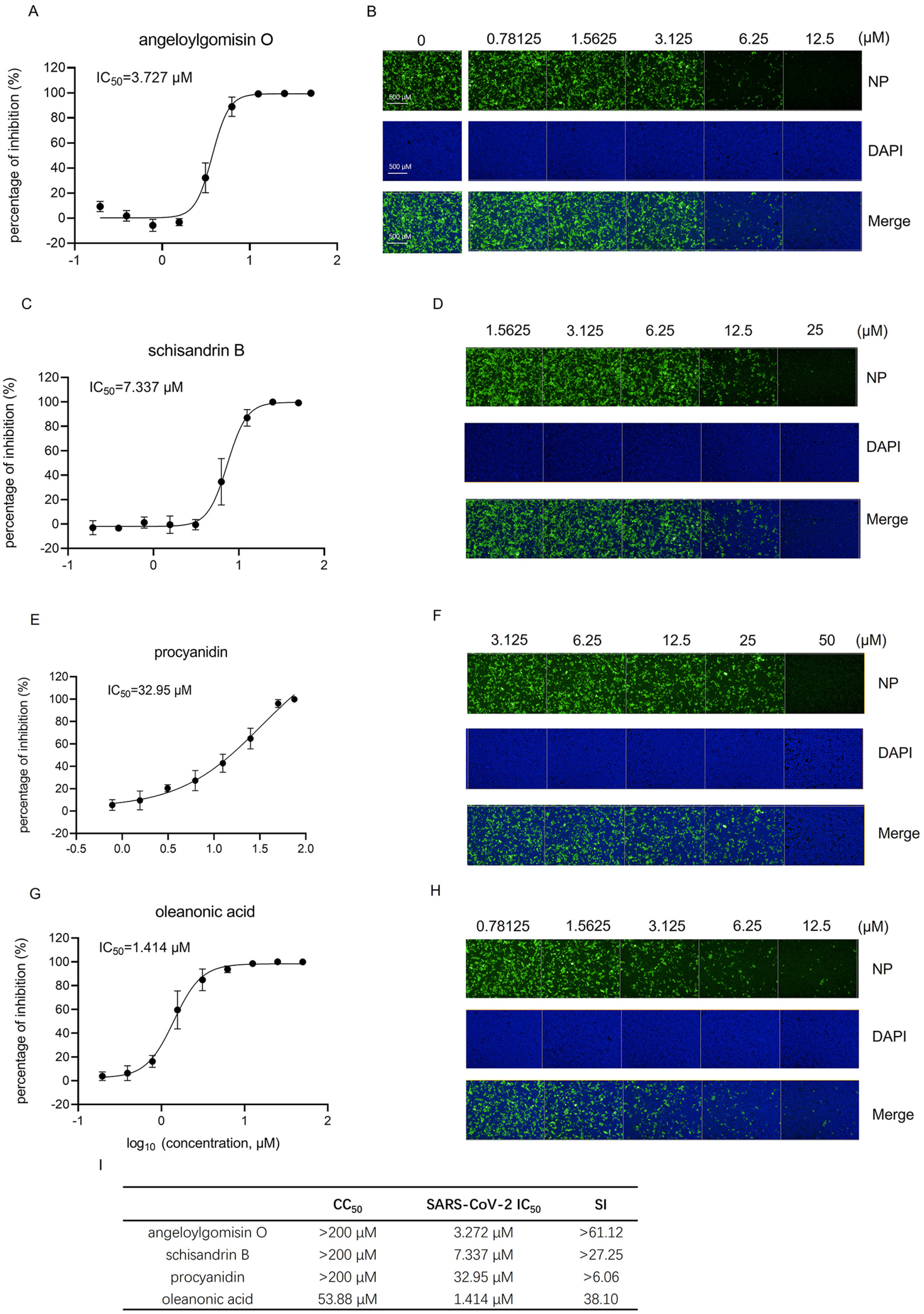
The four hits inhibit authentic SARS-CoV-2 infection. IFA analysis and images of angeloylgomisin O (A and B), schisandrin B (C and D), procyanidin (E and F), and oleanonic acid (G and H) inhibition of SARS-CoV-2 infection in Caco-2 cells. Caco-2 cells were seeded at 96-well plates. After overnight incubation, cell monolayers were treated in duplicate with the hits at a final concentration of 1 h and infected with SARS-CoV-2 at an MOI of 0.5, followed by incubation for 24 h. IFA images showing the viral NP (green) and cell nuclei (blue) are displayed for Caco-2 cells. Cells were treated with different concentrations of angeloylgomisin O (B), schisandrin B (D), procyanidin (F), and oleanonic acid (H). (I) CC_50_, IC_50_, and SI values of four hits. Data were presented as means±SD from three independent experiments.

The selective index (SI, CC_50_/IC_50_) was calculated using the authentic SARS-CoV-2 inhibition assay. The SI values of angeloylgomisin O and schisandrin were > 61.12 and > 27.25, respectively. Suggesting that both drugs are promising potential candidates against COVID-19.

### Effects of the four hits against MERS-CoV-S pv and SARS-CoV-2 S variants pv

To verify the inhibitory effects of the four hits against other coronaviruses and the emergence of SARS-CoV-2 variants, we investigated the antiviral effect of the four hits on MERS-CoV S pv and SARS-CoV-2 S pv variants containing S protein mutations. As shown in Fig. 6A, all the four hits suppressed the entry of MERS-CoV S pv into Caco-2 cells in a dose-dependent manner. The IC_50_ values of angeloylgomisin O, schisandrin B, procyanidin, and oleanonic acid against MERS-CoV S pv were 21.13 μM, 18.99 μM, 13.97 μM, and 18.05 μM, respectively. Indicating that their therapeutic potential against MERS-CoV. MERS-CoV employed the different receptor, dipeptidyl peptidase-4 (DPP4), from the SARS-CoV-2. It could be ruled out that four hits only interfere with ACE2 to block pseudovirus entry (25, 26). A SARS-CoV-2 S pv variant containing the S protein D614G substitution was constructed and used to evaluate its inhibitory effect. As shown in Fig. 6B, all the four hits suppressed the entry of SARS-CoV-2 S pv D614G into Caco-2 cells in a dose-dependent manner. The IC_50_ values of angeloylgomisin O, schisandrin B, procyanidin, and oleanonic acid at which SARS-CoV-2 S pv D614G was inhibited were 16.35 μM, 26.60 μM, 33.36 μM, and 25.21 μM.

**Fig. 6.**
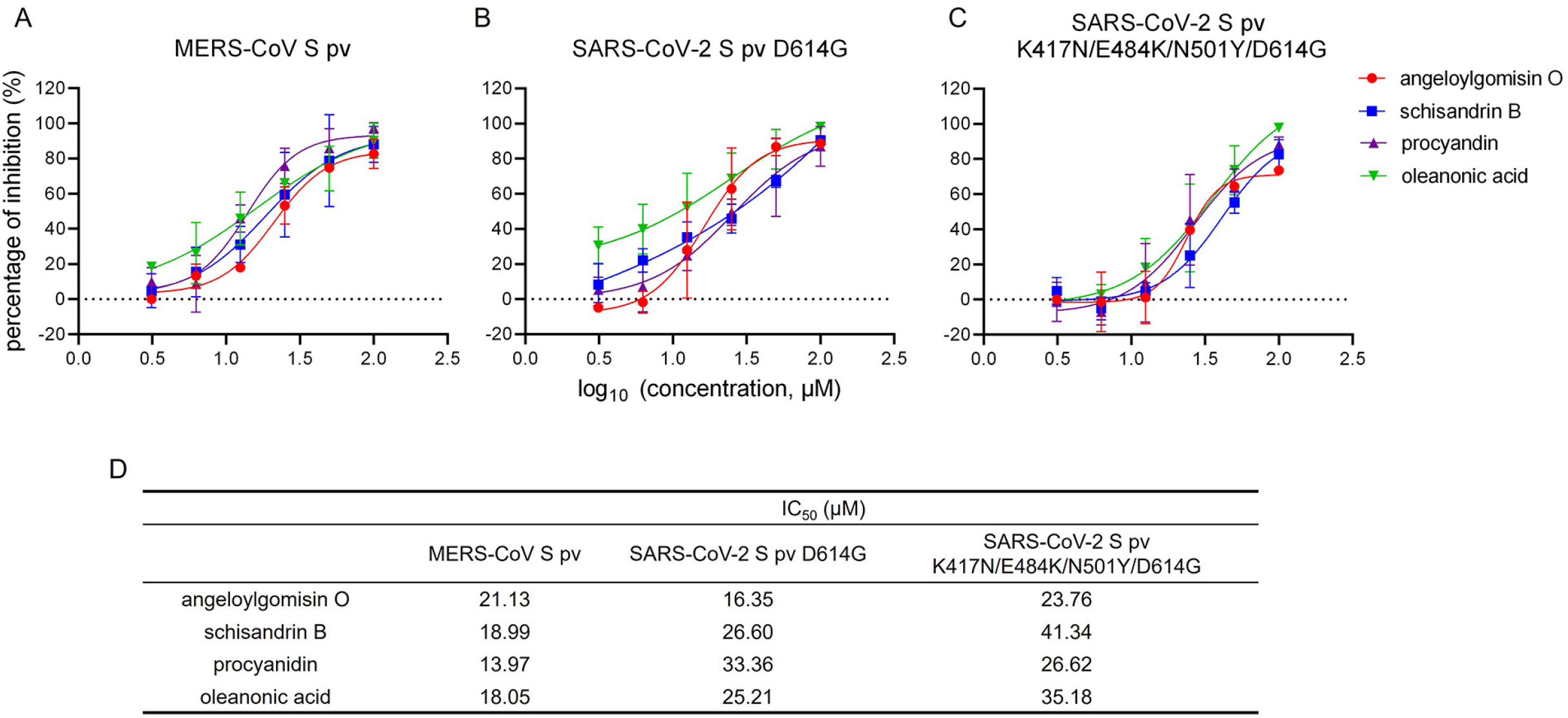
Antiviral activity of the four hits against MERS-CoV S pv (A), SARS-CoV-2 S pv D614G mutant (B), and SARS-CoV-2 S pvK417N/E484K/N501Y/D614G mutant (C). The IC_50_ of four hits for inhibition against the variants MERS-CoV-S pv, SARS-CoV-2 S pv D614G and SARS-CoV-2 S pv K417N/E484K/N501Y/D614G (D). Caco-2 cells were seeded at a density of 2.4 × 10^4^ cells per well in 96-well plates. After overnight incubation, cells were treated in duplicate with the hits at a final concentration; 1 h later, cells were infected with MERS-CoV-S pv, SARS-CoV-2 S pv D614G, and SARS-CoV-2 S pv K417N/E484K/N501Y/D614G with the MOI=5. Data were presented as means±SD from three independent experiments.

We introduced three additional amino acid substitutions, K417N, E484K, and N501Y, in the S D614G protein. These mutations have been observed in the South African variant, 501Y.V2, and play an essential role in immune escape (13–15). A substitution was constructed to evaluate the inhibitory effect. The results showed that all the four hits inhibited SARS-CoV-2 S pv K417N/E484K/N501Y/D614G in a dose-dependent manner (Fig. 6C). The IC_50_ values of angeloylgomisin O, schisandrin B, procyanidin, and oleanonic acid against SARS-CoV-2 S pv K417N/E484K/N501Y/D614G were 23.76 μM, 41.34 μM, 26.62 μM, and 35.18 μM, respectively, indicating that the all the four hits may inhibit new variants such as 501Y.V2. Mutation of K417N/E484K/N501Y on the viral S proteins of 501.V2 variants renders a change in the RBD allowing additional interaction with human ACE2. All the four hits inhibited SARS-CoV-2 S variant with the similar effect. Further confirmed that four hits inhibited membrane fusion rather than interacted with RBD or ACE2 by inhibiting SARS-CoV-2.

### Combinatory treatments with the drug pairs remdesivir-angeloylgomisin O showed enhanced antiviral activity

As all the four hits could inhibit S-mediated membrane fusion, we assessed the efficacy of the combined treatment by using remdesivir, a SARS-CoV-2 viral RNA-dependent RNA polymerase inhibitor (27, 28), and four hits respectively. The degree of interaction was determined according to the bliss independence model using MacSynergy II software to analyze the results (29, 30). Volumes of statistically significant synergy were evaluated, and combination volumes (CV) of > 25 μM^2^ % were interpreted as evidence of synergy, and > 100 μM^2^ %, as strong synergy (31).

Our results showed that strong synergy interactions occurred between remdesivir and angeloylgomisin O. Combination studies between remdesivir and angeloylgomisin O led to the CV of 158.27 μM^2^ % (Fig. 7A). For combinations of remdesivir-angeloylgomisin O the maximal synergistic effect was seen at concentration of 0.032 to 0.8 μM remdesivir and 1.5625 to 6.25 μM angeloylgomisin O. Remdesivir and schisandrin B showed minor amount of synergy CV was 25.3 μM^2^ % (Fig. 7A and 7C). Two-drug combinations of procyanidin-remdesivir and oleanonic acid-remdesivir had an additive antiviral inhibition profile (Fig. 7D and 7E).

**Fig. 7.**
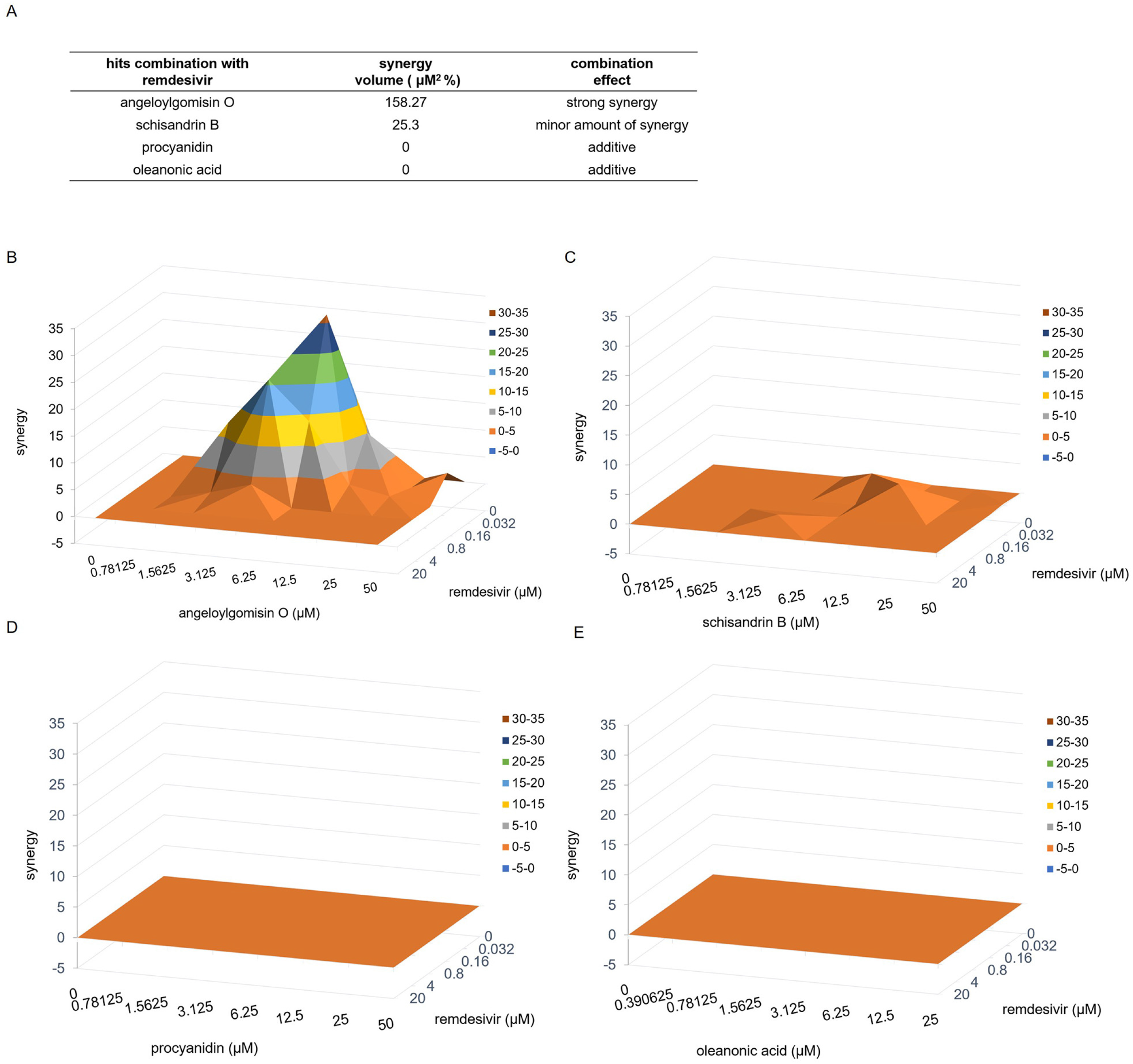
Combinatory treatments with the drug pairs remdesivir-angeloylgomisin O and remdesivir-schisandrin B show enhanced antiviral activity (A) Synergy volumes for pairwise combination studies performed between remdesivir and each hit. Three-dimensional plot of synergy and antagonism at 99% confidence for the pairwise combination of remdesivir with angeloylgomisin O (B), schisandrin B (C), procyanidin (D), oleanonic acid (E). Analysis was performed with MacSynergy II software.

## DISCUSSION

In this study, we screened a botanical drug library containing 1,037 compounds and identified four hits, including angeloylgomisin O, procyanidin, schisandrin B, and oleanonic acid, which blocked the entry of SARS-CoV-2 S (ct19) pv infection by inhibiting viral membrane fusion. Furthermore, the four hits inhibited the SARS-CoV-2 authentic virus at the micromolar level. The top two compounds angeloylgomisin O and schisandrin B inhibited authentic SARS-CoV-2 at a highly selective index (SI). All the four hits inhibited MERS-CoV S pv, SARS-CoV-2 S D614G pv, and SARS-CoV-2 S pv K417N/E484K/N501Y/D614G infection. Remdesivir-angeloylgomisin O drug combination showed stronger antiviral activities against SARS-CoV-2 than remdesivir monotherapy.

The top two hits, angeloylgomisin O and schisandrin B, were derived from *Schisandra chinensis.* Fruit from this plant and its extracts were used in traditional medicine in East Asia to treat liver disorders such as hepatitis. They also possess a broad spectrum of biological and pharmacological uses, including antiviral, anti-inflammatory, and antioxidative properties, without toxicity (32). Schisandrin B has been shown to inhibit cytochrome P450 3A (CYP3A) activity in rat liver and thus affects the metabolism of many drugs (33, 34). Furthermore, upon rat oral administration of *Schisandra chinensis* extract, schisandrin B has been reported to accumulate to a maximum plasma concentration (*C_max_*) of 4.54 μM (35). This value was similar to the IC_50_ value for schisandrin B in the present study, suggesting that this compound offers a biologically feasible treatment for SARS-CoV-2.

At the cellular level, CD4+T cells were rapidly activated to produce inflammatory cytokines after SARS-CoV-2 infection, which further induced CD14+ CD16+ monocyte activation with high levels of expression of interleukin 6 (IL-6). Thus, reduced IL-6 would potentially reduce immunopathology caused by SARS-CoV-2 (36, 37). Cai et al. demonstrated that schisandrin B pretreatment mice at doses of 25, 50, and 75 mg/kg were shown to reduce lipopolysaccharide-induced acute lung injury with lowered the number of inflammatory cells and pro-inflammatory cytokines including tumor necrosis IL-6 in bronchoalveolar lavage fluid (38). This result suggested that schisandrin B could coordinate cytokine response to reduced host immune response, and schisandrin B might achieved a better therapeutic effect in *vivo*.

Procyanidin is a proanthocyanidin member that belongs to the class of flavonoids, the secondary metabolites of polyphenolic plants and fungi. Proanthocyanidins are also the most abundant polyphenolic compounds in lignin (39). Maroli et al. used molecular docking analyses to demonstrate that procyanidin can inhibit SARS-CoV-2 entry and replication (40). In this study, we confirmed that procyanidin inhibits SARS-CoV-2 infection in the entry step.

## MATERIALS AND METHODS

### Cells and viruses

HEK 293T, Vero E6, Caco-2, and BHK-21 cells were cultured in Dulbecco’s modified Eagle medium (HyClone, Logan, UT, USA) supplemented with 10% fetal bovine serum (Gibco, Grand Island, NY, USA). The SARS-CoV-2 strain used in this study is a clinical strain of SARS-CoV-2 (nCoV-2019BetaCoV/Wuhan/WIV04/2019). The pseudotype VSV bearing the S protein of SARS-CoV-2 S (GenBank QHD43416.1) and MERS-CoV S (GenBankNC_019843.3) were generated as previously reported (41, 42). Briefly, 293T cells transfected with pcDNA3.1-SARS-CoV-2 S (ct19), pcDNA3.1-SARS-CoV-2 S (wt), or pcDNA3.1-MERS-CoV S (ct16) for 48 h were infected with pseudotype VSV (described below), in which the G gene was replaced with the luciferase gene, at an MOI of 0.1 for 2 h. The culture supernatants were harvested 24 h later, centrifuged to remove cell debris, and stored at −80°C. To generate VSVG pseudotype VSV, BHK-21 cells in 6-well plates were infected with a recombinant vaccinia virus (vTF7-3) encoding T7 RNA polymerase at an MOI of 5. After 45 min, the cells were transfected with 11 μg of mixed plasmids with a 5:3:5:8:1 ratio of pVSVΔG-eGFP-GPC (pVSVΔG-Rluc to generate pseudotype VSV), pBS-N, pBS-P, pBS-G, and pBS-L. After 48 h, the supernatants were filtered to remove the vaccinia virus and inoculated into BHK-21 cells that had been transfected with pCAGGS-VSV G 24 h previously. The titer of the pseudotype virus was measured by infecting BHK-21 cells previously transfected with pCAGGS-VSV G and determined by plaque assay 24 hours post-infection (h.p.i). The titer of SARS-CoV-2 S pv(ct19) was 1.5×10^7^ PFU/ml.

All point mutants (D614G, K417N/E484K/N501Y/D614G) were made from pcDNA3.1-SARS-CoV-2 S (ct19) by using a fast mutagenesis system (Transgene Biotech), and mutations were confirmed by sequencing (Sangon, Shanghai). We used the same method to generate the SARS-CoV-2 S variants pv.

### Viral copies assay

Equal volumes of SARS-CoV-2 S pv (ct19) and SARS-CoV-2 S pv (wt) were treated with DNase and RNase. RNA from the viruses was extracted using Trizol (TaKaRa) and reverse transcribed using the PrimeScript™ RT reagent Kit (TaKaRa). Viral particles were quantified via RT-qPCR using a specific primer pair to detect VSVΔG-Rluc (primers 5ʹ-GTAACGGACGAATGTCTCATAA-3ʹ and 5ʹ-TTTGACTCTCGCCTGATTGTAC-3ʹ).

### Immunoblotting

Equal volumes of virus particles were precipitated with acetone and lysed using RIPA lysis buffer. Lysates were treated with loading buffer, subjected to SDS-PAGE, and then transferred onto a polyvinylidene difluoride (PVDF) membrane (Millipore). SARS-CoV-2 S was detected using a rabbit anti-S2 subunit mouse monoclonal antibody (GeneTex, GTX632604;1:2000 dilution). Horseraddish peroxidase-linked goat anti-mouse IgG antibody (Proteintech, CHI, USA, 1:5000).

### HTS optimization and assaying of the botanical drug library

The appropriate cell density and the dose for SARS-CoV-2 S pv were selected by comparing the signal-to-basal ratio, the coefficients of variation, and Z’ values under different conditions, as previously described. A library of 1,037 botanical compounds was purchased from Weikeqi Biotech (Sichuan, China). The compounds were collected and stored in 10 mM stock solutions in DMSO at −80 °C until use. The first round of HTS was carried out, and Caco-2 cells were seeded at a density of 2.4×10^4^ cells per well in 96-well plates. After incubation overnight, cells were treated in duplicate with the compounds (50 μM), and 1 h later, cells were infected with SARS-CoV-2 S pv (ct19) with the MOI=5, and the supernatant was removed 1 h.p.i. Camostat 100 μM and 0.5% DMSO were used as positive and negative controls, respectively. After 24 h, luciferase activity was measured using the Rluc assay system (Promega, Madison, WI, USA). The primary compounds were then secondarily screened using VSV pv (MOI of 0.5) to rule out VSV genome replication inhibitors and Rluc activity.

### Cell viability

The 3-(4,5-dimethyl-2-thiazolyl)-2,5-diphenyl-2H-tetrazolium bromide (MTT) assay was performed to evaluate the effect of the four hits on cell viability. Hits at the indicated concentrations were added to pre-seeded Caco-2 cells in 96-well plates. Twenty-four hours later, the cell viability was measured using the MTT kit.

### Membrane fusion assay

Vero E6 cells were co-transfected with pcDNA3.1-SARS-CoV-2S (ct19) (0.25 μg) and pEGFP-N1 (0.25 μg) by using lipo2000 in 24-well plates. After transfection for 4 h, the medium was replaced with 2% DMEM containing different concentrations of the hits. After 24 h, syncytium formation was visualized using an M-shot image system (Micro-shot Technology, China)

### Antiviral effect of four hits against SARS-CoV-2

Caco-2 cells were seeded at a density of 2.4×10^4^ cells per well in 96-well plates. After overnight incubation, cell monolayers were treated in duplicate with the hits at a final concentration of 1 h and infected with SARS-CoV-2 at an MOI of 0.5. After an additional 24 h of incubation, the infection was stopped by rinsing each well, and the cells were fixed with 4% paraformaldehyde. Cells were permeabilized using PBS with 0.2% Triton X-100 for 15 min and blocked with 5% FBS (Gibco), followed by treatment with the primary antibody anti-SARS-CoV-2 NP (GeneTex GTX635678, USA) at a 1:500 dilution overnight at 4°C. After six rinses with PBS, the cells were stained with DyLight 488-labeled antibody against rabbit IgG (KPL, Gaithersburg, MD, USA). Nuclei were stained with 4’,6-diamidino-2-phenylindole (DAPI, Sigma-Aldrich, St. Louis, MO, USA). Nine fields per well were imaged, and the percentages of infected and DAPI-positive cells were calculated using Harmony 3.5 software.

### Drug-drug interactions of remdesivir with hits

Caco-2 cells were seeded at a density of 2.4×10^4^ cells per well in 96-well plates. After overnight incubation, different final concentrations of remdesivir were added to each row of the 96-well plate. Simultaneously, different final concentrations of angeloylgomisin O were added to each column of the plate. The cells were infected with SARS-CoV-2 at an MOI of 0.5. After an additional 24 h of incubation, the infection was stopped by rinsing each well, and the cells were fixed with 4% paraformaldehyde. The antiviral activities were determined using the IFA assay. To determine the drug-drug interactions, differential surface plots at the 95% confidence level (CI) were generated using MacSynergy II according to the Bliss independence model. The combinatorial effect was determined by subtracting the theoretical additive values from the experimental values (29, 30). The data sets were interpreted using the following scale: volumes of synergy or antagonism > 100 μM ^2^ %, strong synergy; 50–100 μM ^2^ %, moderate synergy; 25–50 μM ^2^ %, minor amount of synergy; 0 −25 μM ^2^ %, additive (31).

## ACKNOWLEDGMENT

We thank Tao Du and Jin Xiong from the BSL-3 Laboratory of the Wuhan Institute of Virology for their critical support. We thank the Center for Instrumental Analysis and Metrology, Core Facility, and Technical Support, Wuhan Institute of Virology, for providing technical assistance.

This work was supported by the National Key Research and Development Program of China (2018YFA0507204), the National Natural Sciences Foundation of China (31670165), Wuhan National Biosafety Laboratory, Chinese Academy of Sciences Advanced Customer Cultivation Project (2019ACCP-MS03), and the Open Research Fund Program of the State Key Laboratory of Virology of China (2018IOV001).

